# Aggregation chimeras provide evidence of *in vivo* intercellular correction in ovine CLN6 neuronal ceroid lipofuscinosis (Batten disease)

**DOI:** 10.1101/2021.12.06.471461

**Authors:** Lucy A. Barry, Graham W. Kay, Nadia L. Mitchell, Samantha J. Murray, Nigel P. Jay, David N. Palmer

## Abstract

The neuronal ceroid lipofuscinoses (NCLs; Batten disease) are fatal, mainly childhood, inherited neurodegenerative lysosomal storage diseases. Sheep affected with a CLN6 form display progressive regionally defined glial activation and subsequent neurodegeneration, indicating that neuroinflammation may be causative of pathogenesis. In this study, aggregation chimeras were generated from homozygous unaffected normal and CLN6 affected sheep embryos, resulting in seven chimeric animals with varied proportions of normal to affected cells. These sheep were classified as affected-like, recovering-like or normal-like, based on their cell-genotype ratios and their clinical and neuropathological profiles.

Neuropathological examination of the affected-like animals revealed intense glial activation, prominent storage body accumulation and severe neurodegeneration within all cortical brain regions, along with vision loss and decreasing intracranial volumes and cortical thicknesses consistent with ovine CLN6 disease. In contrast, intercellular communication affecting pathology was evident at both the gross and histological level in the normal-like and recovering-like chimeras, resulting in a lack of glial activation and rare storage body accumulation in only a few cells. Initial intracranial volumes of the recovering-like chimeras were below normal but progressively recovered to about normal by two years of age. All had normal cortical thicknesses, and none went blind. Extended neurogenesis was evident in the brains of all the chimeras.

This study indicates that although CLN6 is a membrane bound protein, the consequent defect is not cell intrinsic. The lack of glial activation and inflammatory responses in the normal-like and recovering-like chimeras indicate that newly generated cells are borne into a microenvironment conducive to maturation and survival.

## Introduction

The neuronal ceroid lipofuscinoses (NCLs; Batten disease) are fatal lysosomal storage diseases that collectively constitute one of the most common childhood inherited neurodegenerative diseases. They also occur in animals. Different forms are caused by mutations in any one of 13 different genes, designated *CLNs 1-8* and *10-14* ([1], http://www.ucl.ac.uk/ncl). The group is defined by the near ubiquitous accumulation of protein, either the c subunit of mitochondrial ATP synthase or the sphingolipid activator proteins (SAPs) A and D, in lysosome-derived storage bodies in neurons and most somatic cells [1–4]. The other defining feature of the NCLs is regionally specific neurodegeneration and brain atrophy [5]. Clinical features are progressive mental and motor deterioration, blindness and seizures leading to premature death, usually between 7 years of age and early adulthood [1, 2].

The best characterised animal model of the disease is a CLN6 form in New Zealand South Hampshire sheep in which disease progression closely parallels that of variant late infantile human CLN6 [6–9]. CLN6 affected sheep develop clinical symptoms between 10 and 14 months, namely loss of vision and progressive neurological decline, and die prematurely, usually before 24 months [6]. Regionally defined and selective loss of cortical neurons in the CLN6 affected sheep is preceded by prenatal neuroinflammation, beginning in particular cortical foci which are associated with later symptomology [10–13].

Understanding the interconnections between the genetic lesion, lysosomal storage, and neurodegeneration is pivotal for determining options for therapy. *CLN6* encodes an endoplasmic reticulum resident protein of uncertain function [14–16], thus CLN6 disease is usually regarded as a cell intrinsic disorder unlikely to benefit from a therapeutic strategy reliant on cross-correction, whereby soluble lysosomal proteins secreted from cells are endocytosed into protein-deficient cells [17, 18]. However the cellular loss in affected sheep is primarily restricted to the central nervous system (CNS) despite widespread storage body accumulation in most somatic cells, suggesting that location and connectivity, not phenotype or storage burden, determine neuronal survival in ovine CLN6 [3, 12]. Furthermore, there is much evidence in the literature to suggest that some degree of intercellular transport and cross-correction could occur in CLN6 disease. For example, it is possible if the membrane bound CLN6 processes a soluble factor, perhaps in a way similar to reports that CLN7 processes CLN5 [19] or as part of a CLN6-CLN8 complex that recruits lysosomal enzymes at the ER for Golgi transfer [20], or if it modulates the expression of other glycosylated lysosomal hydrolases as suggested in CLN6 affected *nclf* mouse studies [21].

The generation of chimeric animal models provides a direct way to test whether affected cells are amenable to correction by unaffected normal cells *in vivo* [22–25]. For this study, aggregation chimeras were generated from homozygous unaffected normal and CLN6 affected sheep embryos. The resultant chimeras possessed varying proportions of normal and affected cells and clinical and neuropathological profiles somewhere between those of affected and normal animals. Factors analysed in the chimeras included cortical atrophy, a definitive hallmark of NCL, neuroinflammation that has been strongly implicated in disease pathogenesis [11, 13] and evidence of extended neurogenesis and clusters of newly generated neurons in the affected brain [26].

## Materials and methods

### Animals

Sheep were maintained under standard New Zealand pastoral conditions on Lincoln University farms. Animal procedures were approved by the Lincoln University Animal Ethics Committee (LUAEC#213) and accorded with the New Zealand Animal Welfare Act (1999) and US National Institutes of Health guidelines. Black faced homozygous CLN6 affected South Hampshire sheep were bred and diagnosed using a discriminatory c.822 G>A single nucleotide polymorphism (SNP) in ovine *CLN6* [9]. White faced homozygous unaffected Coopworth sheep were used as normal controls.

### Production of aggregation chimeras

Normal and affected ewes were synchronised with progesterone-impregnated controlled intrauterine drug release devices, induced to superovulate using follicle stimulating hormone, then fertilised by laparoscopic insemination [27]. Embryos at the 16-32 cell morula stage were collected by flushing the uterine horns and approximately half the blastomeres from selected homozygous affected embryos were exchanged for approximately half the blastomeres of selected homozygous normal embryos (Fig 1a). The resultant hybrid embryos were re-implanted into synchronised normal recipient ewes for development to term, yielding 15 lambs.

**Fig 1.**
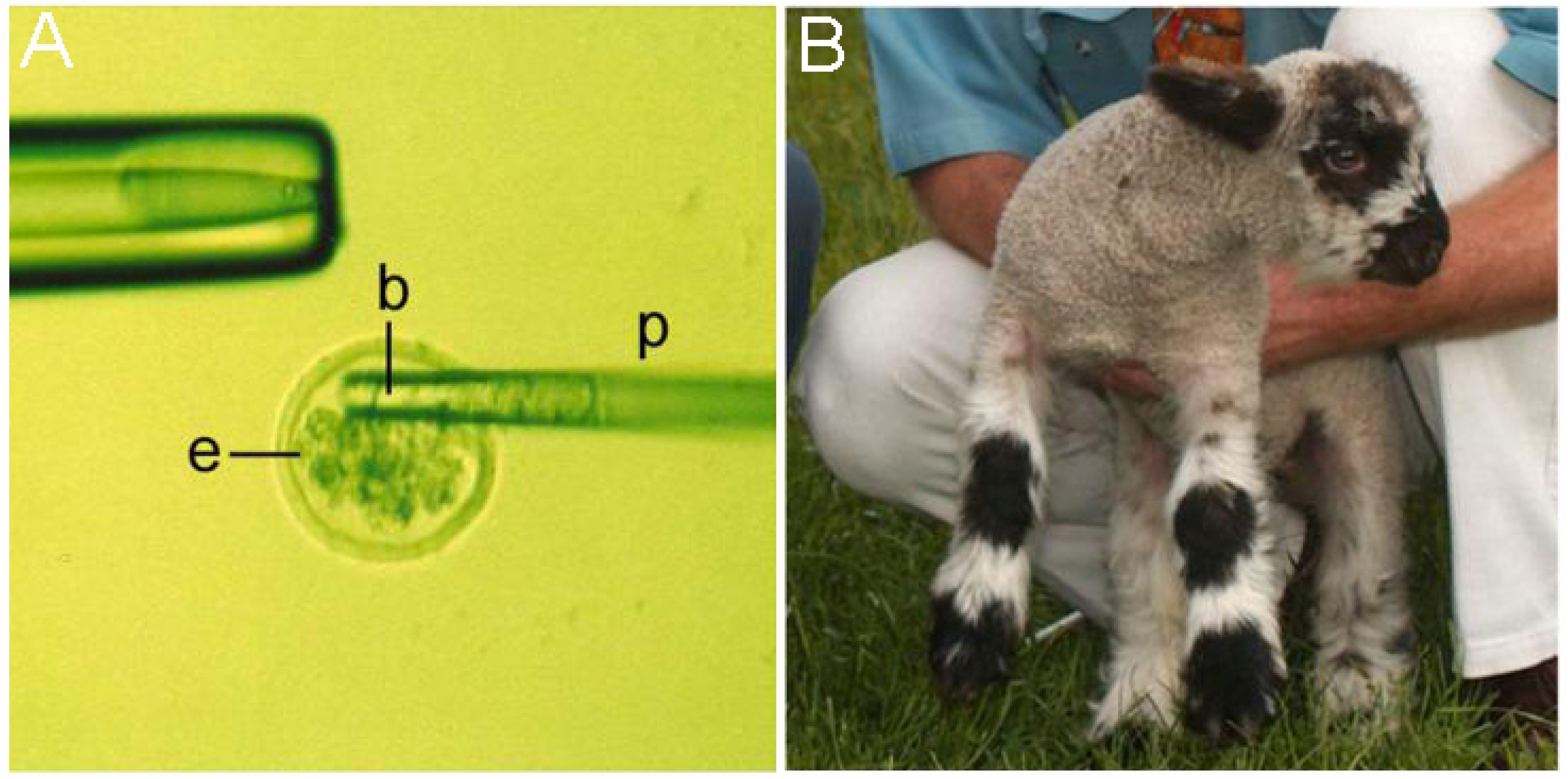
Exchange of blastomeres between homozygous normal and CLN6 affected embryos to create chimeric lambs. **A)** Blastomeres (b) of an affected embryo positioned in the tip of an aspiration pipette (p) being deposited into a normal embryo (e) after removal of approximately half of the normal blastomeres. **B)** A resultant chimeric lamb displaying a distinctive chimeric coat pattern.

### Development of chimeric lambs

Computed tomography (CT) scans of these lambs were first performed between 2-6 months of age and then approximately every 6 months thereafter. They were compared to historical data from affected (n=43 scans) and normal (n=54 scans) controls from the same flocks. Coronal slices, 1 mm thick, were obtained at 5 mm intervals, 120 KV, 160 ma, 2 s, in a GE Pro-speed CT scanner (GE Healthcare, Hyogo, Japan) and intracranial volumes were determined by the Cavalieri method from the areas of each slice [28] to estimate brain volumes [29]. Body weights were also compared to those of normal and affected animals, and clinical loss of vision assessed by a simple obstacle course test and a blink response to bright light [30].

### Tissue collection and processing

Chimeric sheep were euthanised by exsanguination between the ages of 17 and 41 months, when severe clinical disease symptoms became apparent, or it was evident that no clinical symptoms were developing. Two CLN6 affected and two normal Coopworth animals, aged 18 and 24 months, served as controls. Brains were removed intact, halved down the midline and immersion fixed in 10% formalin in 0.9% NaCl. Tissue samples of endodermal (liver, thyroid, pancreas), mesodermal (cardiac and skeletal muscle, kidney, testis, ovary) and ectodermal (brain, skin) embryonic germ layer origin were also immersion fixed or snap frozen in liquid nitrogen and stored at -80°C. Eyes were enucleated and immersed in 10% formalin, then sent to Gribbles Veterinary pathologists (Christchurch, New Zealand) for post-fixation in Bouin’s solution and wax embedding.

The fixed brains were equilibrated, frozen at -80°C, then sectioned in the sagittal plane at 50 μm [31]. Matched mediolateral level 4 sections were selected from each brain for subsequent immunohistochemical analyses [13].

### Immunohistochemical staining and histology

All antibodies were diluted in 10% normal goat serum in phosphate buffered saline (PBS), pH 7.4, containing 0.3% Triton X-100 (Sigma Aldrich, St Louis, MO, USA). Primary antibodies used were rabbit anti-cow glial fibrillary acidic protein (GFAP; 1:5000; Z0334; Dako, Carpinteria, CA, USA) to detect astrocytes, a biotinylated form of the α-D-galactose specific isolectin I-B4 from *Griffonia simplicifolia* (GSB4; 1:500; B-1205; Vector Laboratories, Burlingame, CA, USA; 1:500) for activated microglia detection, and mouse anti-polysialated neural cell adhesion molecule (PSA-NCAM; 1:1000; MAB5324; Chemicon, Temecula, CA, USA) for detecting newly generated and migrating cells.

Immunohistochemical detection used the avidin-biotin amplification system [11–13], appropriate secondary antibodies and ExtrAvidin peroxidase (1:1000; B7389; Sigma Aldrich). The optimal incubation period with 3, 3’-diaminobenzadine substrate solution (0.5mg/ml; D5637, Sigma Aldrich) in 0.01% H_2_O_2_ in PBS was determined for each antigen. Negative control sections with either the primary or secondary antibody omitted were included in all staining runs. Sections were mounted in a solution of 0.5% gelatine and 0.05% chromium potassium sulphate on glass slides, air-dried, dehydrated in 100% ethanol, cleared in xylene and coverslipped with DPX (BDH, Poole, England).

Nissl and Luxol fast blue staining was performed on mounted sagittal brain sections as described [13]. Corresponding sets of unstained sections were mounted and coverslipped with glycerol for analysis of storage body fluorescence.

Retinal paraffin sections were cut at 3 µm, mounted and a subset stained with hematoxylin and eosin (H+E) histological stain by Gribbles Veterinary Pathology for analysis of retinal thickness. Unstained retinal sections were also coverslipped in glycerol for assessment of of storage body fluorescence.

### Microscopy and image analysis

Digital images of stained sections were acquired with an inverted DMIRB microscope (Leica, Wetzlar, Germany) using a SPOT RT colour digital camera and software (v4.0.9, Diagnostic Instruments Inc., Sterling Heights, MI, USA). Lamp intensity, exposure time, condenser aperture setting, video camera setup and calibration, and the use of a neutral density filter was kept constant for capturing all images of a particular immunostain. Digital images were saved as .tif files and figures and photomontages prepared in Corel Photopaint 12 (Corel Co., Ontario, Canada).

Fluorescent storage body accumulation was determined by confocal laser scanning microscopy of unstained sections (Leica, TCS SP5) using 405 nm excitation and 535 nm emission filters. Images were obtained from both cortical and cerebellar regions. The pinhole size, amplitude off-set and detector gain settings were kept constant for all sections. Confocal images (.lsm) were converted to .tif files using LAS AF lite software (Leica) and figures prepared in Corel Photopaint 12. All sections were analysed independently by a second person without knowledge of the genotype. All observations were consistent to both viewers.

GFAP and GSB4 stained sections were imaged under bright-field microscopy and thresholding analysis of images was performed on ImageJ (NIH, version 1.52P) to determine the percentage stained per sampled area.

All processed retinal sections were imaged on a Nikon Eclipse 50i light microscope (Nikon Instruments Inc., Tokyo, Japan) paired to a Nikon Digital Sight DS-U3 camera and NIS-Elements BR software (v. 4.50 Nikon Instruments). Retinal thickness measurements were taken from the surface of the nerve fibre layer (NFL) to the base of the retinal pigment epithelium in the central retina. Fluorescent lysosomal storage was imaged using a GFP Brightline 490 nm excitation/510 nm emission filter set (GFP-3035C; Semrock Inc, IDEX Corporation, IL, USA). Thresholding analysis was performed on ImageJ to determine the percentage of fluorescence per sampled area.

### Grey matter thickness measurements

At least 25 cortical grey matter thickness measurements were made on Nissl stained sections through the occipital cortex as previously described [13].

### DNA and RNA extraction

Genomic DNA (gDNA) was extracted from tissues from chimeric, homozygous CLN6 affected, CLN6 heterozygous and homozygous normal sheep using an Axyprep Multisource Genomic DNA Miniprep Kit (Axygen Scientific Inc, CA, USA). Tissues sampled included blood, liver, thyroid, pancreas, kidney, testis, ovary, skin, cardiac and skeletal muscle, and from several CNS sites; the frontal, parietal and occipital cortex, thalamus, cerebellum, brainstem and the spinal cord. Total RNA was extracted from the same CNS sites using Qiagen RNeasy mini kits (Qiagen, Hilden, Germany). Complementary DNA (cDNA) was generated from 200 ng/µl of total RNA using Superscript III reverse transcriptase (18080044; Invitrogen, Carlsbad, CA, USA) and random hexamer primers.

### PCR genotyping of chimeric animals

An indirect DNA test based on the discriminatory c.822 G>A SNP in *CLN6* exon 7 was used to estimate the ratios of normal and affected cells within tissues [9]. The sheep flocks are structured so that homozygously normal animals (cells) are GG and affected animals (cells) AA, hence the G:A ratio in a tissue is indicative of its chimerism.

PCR reactions were carried out on a Mastercycler Gradient PCR machine (Eppendorf, Hamburg, Germany). Standard 20 µl PCR reactions were performed [9], but at a T_m_ of 55 °C, and included either 80 ng/µl gDNA or 1 µl of cDNA and 0.125 µM of the forward E7F1 (5′-GTA CCT GGT CAC CGA GGG-3′) and reverse 7aR (5′-AGG ACT CTA TTG GCT GC-3′) primers. CLN6 affected, CLN6 heterozygote and normal control gDNA or cDNA was included in all PCR runs. The resulting 277 bp (E7F1-7aR) PCR products were digested with 1 U *Hae*II (NEBR0107S; New England Biolaboratories, Ipswich, MA, USA), 24 h, 37 °C and the fragments separated on 3.5% agarose gels. Normal GG sheep yielded three fragments of 67, 91 and 119 bp; heterozygous GA sheep four fragments of 67, 91, 119 and 186 bp; and CLN6 affected AA sheep two fragments of 91 and 186 bp (Fig 2). Serial dilutions of 80 ng/µl homozygous CLN6 affected gDNA and 80 ng/µl homozygous normal gDNA were then prepared to provide samples with known affected:normal DNA ratios. PCR, *HaeII* restriction digestion and gel electrophoresis were performed as described [9]. Chimeric samples were then compared visually to the serial dilutions for estimations of the affected:normal cell ratios (Fig 2), reported as a % of affected DNA present in each sample.

**Fig 2.**
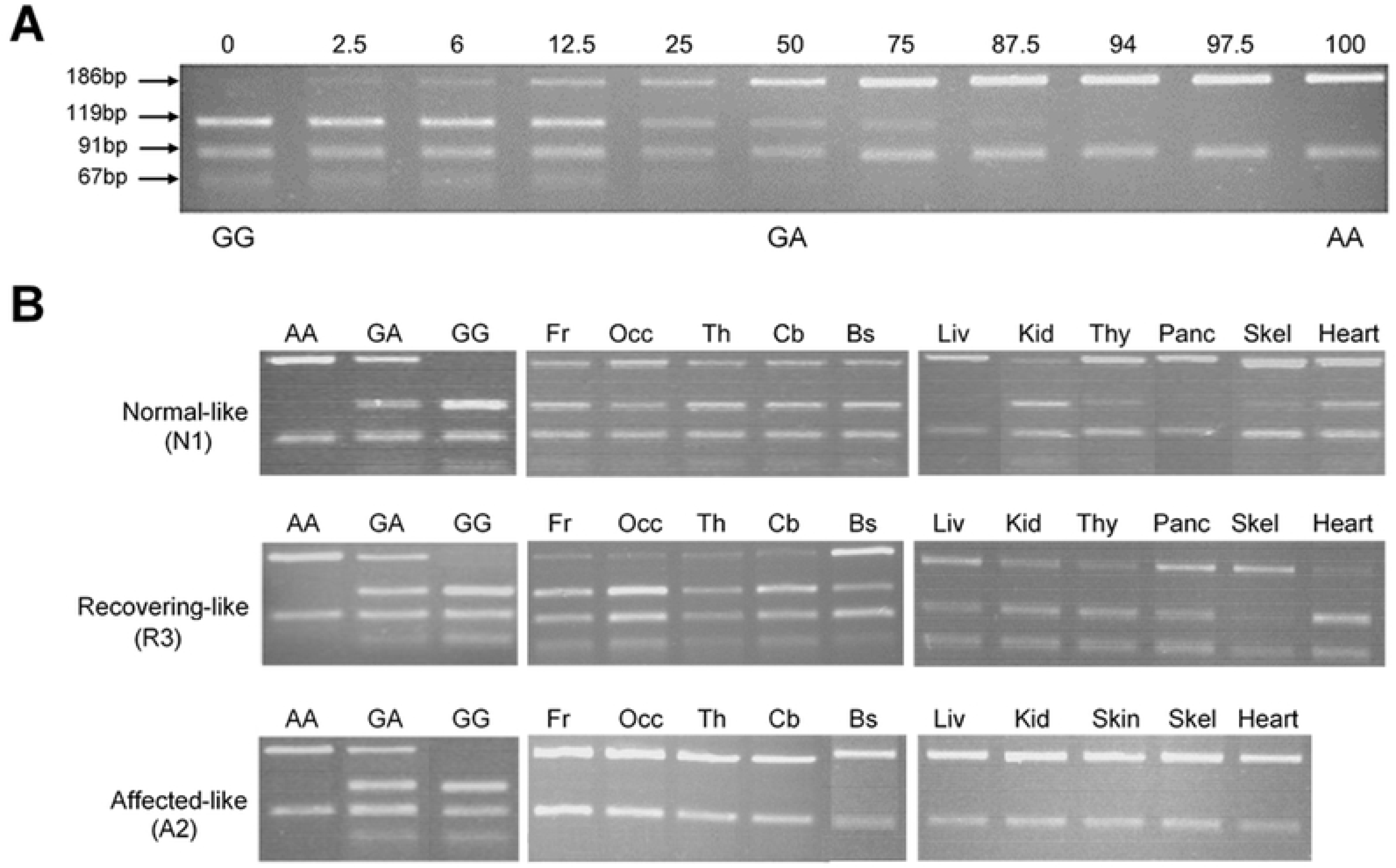
Restriction enzyme detection of the c.822G>A polymorphism to determine the extent of chimerism. A 277bp PCR product from normal **(GG)** sheep cleaved with *HaeII* results in three bands of 119, 91 and 67bp, affected **(AA)** sheep yield two bands of 186 and 91bp, and heterozygous **(GA)** sheep four bands of 186, 119, 91 and 67bp. **A)** Serial dilutions of affected and normal DNA which were used as a standard to estimate the proportion of affected DNA present in samples from chimeric animals by visual inspection. Numbers indicate the affected portion (%). **B)** Representative images of PCR products from brain and peripheral tissues for one animal from each experimental group (See Table 1 for full data set). Fr: Frontal cortex, Occ: Occipital cortex, Th: Thalamus, Cb: Cerebellum, Bs: Brainstem, Sc: Spinal Cord, Liv: Liver, Kid: Kidney, Thy: Thyroid, Panc: Pancreas; Skel: skeletal muscle.

**Table 1.** Genotyping tissues of different embryonic germ layer origins in chimeric animals.

| Animal | Blood | Ectodermal |  |  |  |  |  | Endodermal |  |  | Mesodermal |  |  |
| --- | --- | --- | --- | --- | --- | --- | --- | --- | --- | --- | --- | --- | --- |
|  |  | Frontal lobe | Occipital lobe | Thalamus | Cerebellum | Brainstem | Skin | Liver | Thyroid | Pancreas | Kidney | Skeletal muscle | Cardiac muscle |
| A1 | 100 | NT | NT | NT | 100 | 100 | 25 | 12.5 | 100 | NT | 100 | 25 | 50 |
| A2 | 100 | 100 | 100 | 100 | 100 | 100 | 100 | 100 | NT | NT | 100 | 100 | 100 |
| R1 | 3 | 6 | 25 | 6 | 3 | 12.5 | NT | 0 | 0 | 0 | 0 | 0 | 50 |
| R2 | 0 | 6 | 6 | 3 | 3 | NT | NT | 0 | 0 | 0 | 0 | 0 | 3 |
| R3 | 12.5 | 6 | 6 | 25 | 6 | 75 | NT | 25 | 25 | 25 | 12.5 | 50 | 6 |
| N1 | 0 | 12.5 | 12.5 | 25 | 50 | 12.5 | NT | 87.5 | 75 | 87.5 | 12.5 | 75 | 50 |
| N2 | 50 | 25 | 25 | 3 | 3 | 25 | NT | 50 | 25 | 25 | 6 | 6 | 75 |

Abbreviations: A, affected-like; R, recovering-like; N, normal-like; NT, not tested (sample not available)

### Sequencing

Chimeric status was confirmed via sequencing. PCR products were separated on agarose gels, excised and purified using a CleanSEQ Dye-Terminator removal kit (Agencout Bioscience Corporation, Beverly, MA, USA). Sequence reactions were performed using a BigDye terminator v3.1 Cycle sequencing kit [9, 32] and sequences aligned against the ovine *CLN6* exon 7 sequence (Genbank accession number NM_001040289) [9] to confirm which nucleotide was present at the c.822 G>A SNP site.

### Statistical analysis

All statistical analysis was performed on GraphPad Prism (v 8.2.0, GraphPad Software). Histological analyses were reported as the means ± the standard error of the mean (SEM). Differences between the normal-like (n=2), recovering-like (n=3), and affected-like (n=2) chimeras, and age-matched healthy (n=2) and affected (n=2) controls were assessed using a one-way ANOVA with Tukey’s multiple comparisons test. Where homoscedasticity was not assumed, Brown-Forsythe and Welch ANOVA tests were performed with Dunnett’s T3 multiple comparisons test. Results were considered significant where p<0.05.

## Results

Simplistically the chimera generation strategy of fusing half the blastomeres from homozygous affected embryos and homozygous normal embryos (Fig 1A) should result in 50:50 affected:normal cell chimeras, however the lambs generated here had a wide range of affected:normal cell ratios, which also varied between different tissues within each animal. Contributions to chimerism in each lamb were assessed from coat-colour patterns (Fig 1B), blood DNA analysis and monitoring for any signs of clinical disease. Of the 15 lambs generated during this study, seven were classed as chimeric and kept for further analysis, while six were disregarded as 100% normal and two as 100% affected.

### Genotype analysis and intracranial volume development

The c.822 G>A polymorphism in the ovine *CLN6* exon 7 [9] was exploited for assessment of the ratio of affected:normal cells in a range of tissues from the seven chimeric animals. These results were correlated with intracranial volume data and the chimeric lambs classified into affected-like (A), recovering-like (R) and normal-like (N) groups, defined by the overall whole animal phenotype.

Most tissues from two animals, A1 and A2, were dominated by affected cells, all CNS samples yielding the affected genotype banding pattern (e.g. A2, Fig 2). Blood samples analysed from these animals also correlated with this tissue genotype (Table 1). Their intracranial volume changes closely followed those in affected sheep, decreasing progressively to 70-75 mL by 24 months compared to 110 mL for normal control animals of the same age (Fig 3). Terminal bodyweights of these animals (37.0 and 43.4 kg) were well under half those of normal controls (85-95 kg) and they also lost their vision.

**Fig 3.**
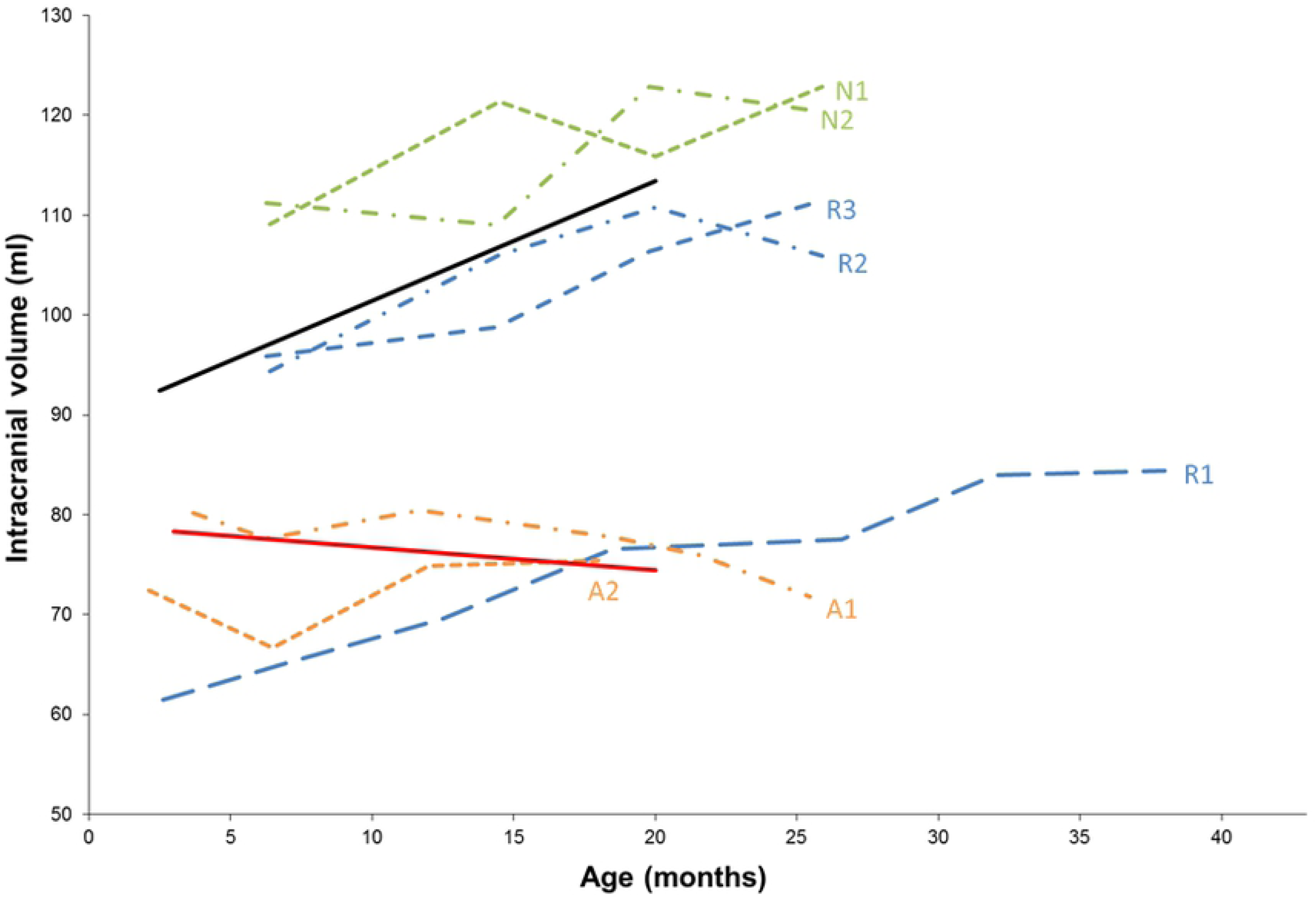
Changes in intracranial volumes. Intracranial volumes of the seven chimeric animals were compared to historic trend data from affected (red line, n=43 scans) and normal (black line, n=54 scans) controls. Green: normal-like (N) animals, Blue: recovering-like (R) animals, Orange: affected-like (A) animals.

The other five chimeric animals; R1, R2, R3, N1 and N2, had both normal and affected cells present in varying proportions within the brain (e.g. R3 and N1, Fig 2). Most brain regions in these animals had a higher proportion of normal than affected cells present but there were some dramatic variations, such as the brainstem of animal R3 which was 75% affected whereas other regions in this brain had more normal cells (Fig 2).

Genotype results from the non-CNS tissues also indicated chimerism, with the percentage of affected cells varying noticeably between tissues of endodermal, mesodermal and ectodermal origin in most animals (Table 1). Analyses of blood samples revealed the presence varying amounts of affected DNA in animals R1, R3 and N2 which was not found in samples from animals N1 and R2 (Table 1). Reverse transcription of RNA with subsequent cDNA amplification and restriction enzyme digestion confirmed that gene presence was reflective of gene expression within a particular tissue.

Animals R2 and R3 had intracranial volumes within the normal range, which progressively increased throughout the study (Fig 3). A 4.9 mL reduction in intracranial volume was observed in animal R2 at 25 months, but volumes were still significantly greater than those of affected animals who rarely survive beyond two years. At baseline (3 months), the brain volume of animal R1 was well below the affected trend-line, 60 mL compared to ∼80 mL for age-matched affected sheep (Fig 3). Its intracranial volume progressively increased, surpassing the affected trend-line at 20 months and continued to increase over the following 20 months to approach the normal line, in contrast to the progressive decrease in intracranial volume seen in affected animals. Of note, the lifespan of these three animals was extended, particularly animal R1 which was not sacrificed until 41 months of age, and they retained healthy body weights (70.2 – 94.5 kg) and did not lose their sight. Based on their intracranial volume data, in combination with the genotype analysis which revealed the presence of affected cells in all brain regions (Table 1), these three animals, R1, R2 and R3, fell into the category of recovering-like.

Some fluctuations in intracranial volume were observed for the remaining two animals, N1 (Fig 2) and N2. Nevertheless, their volumes progressively increased and were consistently above the normal trend-line. They too had healthy terminal body weights (71 – 86 kg) and retained vision. Hence, these two chimeric sheep were classified as normal-like, albeit with some colonisation of affected cells in both CNS and non-CNS tissues as indicated by genotyping (Table 1).

### Cortical atrophy and general organisation of the chimeric brains

Next the *in vivo* data for these seven chimeras were correlated with histochemical examinations of glial activation, neurogenesis, neurodegeneration and storage body accumulation at *post mortem* to determine any influences normal cells might have had on affected cells and on the development of pathology within the brain.

Neurodegeneration and cortical thickness changes were analysed in Nissl stained sections. Marked atrophy of the cerebral cortex and thinning of the cortical layers was discernible in all regions of the two affected-like chimeric animals, A1 and A2 (Fig 4). This was most pronounced in the occipital cortex, where cortical thickness measurements were reduced to 48-52% of normal by 24 months of age, comparable with that of affected animals (Fig 4). Nissl staining revealed widespread neuronal loss in all cortical regions in these animals and a change from a laminar distribution of cells towards densely packed cellular aggregates, particularly at the layer I/II interface of the cortical grey matter (Fig 4).

**Fig 4.**
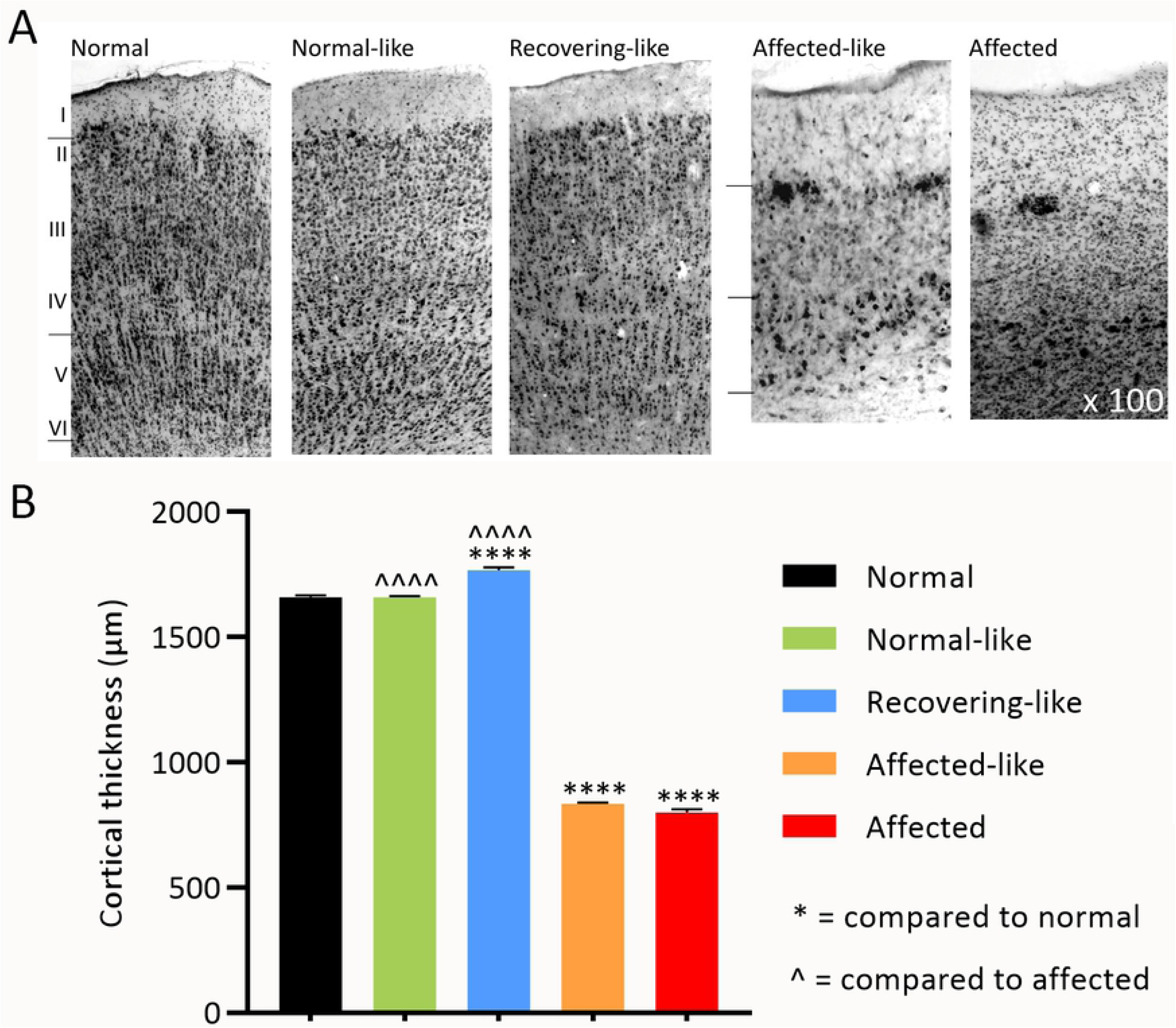
Nissl staining of the occipital cortex. **A)** Representative images of normal-like and recovering-like chimeras show a typical normal cortical layer structure with no indication of neurodegeneration. Extensive neuronal loss and less distinct cortical layer boundaries are evident in affected-like chimeras with densely packed cellular aggregates visible within upper cortical layers. See S1 Fig for all images. **B)** Thickness measurements were made through the occipital cortex of chimeric sheep and compared to adult normal (black) and affected (red) controls. The five recovering-like (blue) and normal-like (green) chimeras are within the normal range whereas affected-like (orange) chimeras have cortical thickness measurements equivalent to affected controls. Note: n ≥ 50 measurements. **** and ^^^^^ indicate P<0.0001.

The two normal-like chimeras, N1 and N2, and the three recovering chimeras, R1, R2 and R3, all displayed a laminar distribution of cells within all cortical regions which closely resembled that in the normal sheep cortex, there being no overt loss of neurons or formation of cellular aggregates (Fig 4), regardless of the degree of chimerism revealed by DNA analysis (Table 1). Cortical thickness measurements quantitated these findings, all normal-like and recovering chimeras being within 97-114% of normal at 24 months of age or trial completion (Fig 4). The cytoarchitecture of the cerebellum and hippocampus remained unchanged in all chimeric animals.

### Storage body accumulation within the chimeric brains

Histological studies revealed the presence of storage bodies in the brains of the affected-like chimeric animals, A1 and A2, consistent with an affected diagnosis (Fig 5). They were fluorescent, stained strongly with Luxol-fast blue and were evident throughout all neocortical regions. The few large pyramidal neurons remaining in the affected-like chimeric cortices were densely packed with globular storage body deposits (Fig 5A) while smaller neuronal and glial-like cells were predominantly filled with granules which occupied the entire cytoplasm. Subcortical and cerebellar regions contained many storage deposits, most obvious within the perikarya of large Purkinje cells (n >200/ cells viewed), the majority of which contained globular, punctate storage bodies of varying size (Fig 5B).

**Fig 5.**
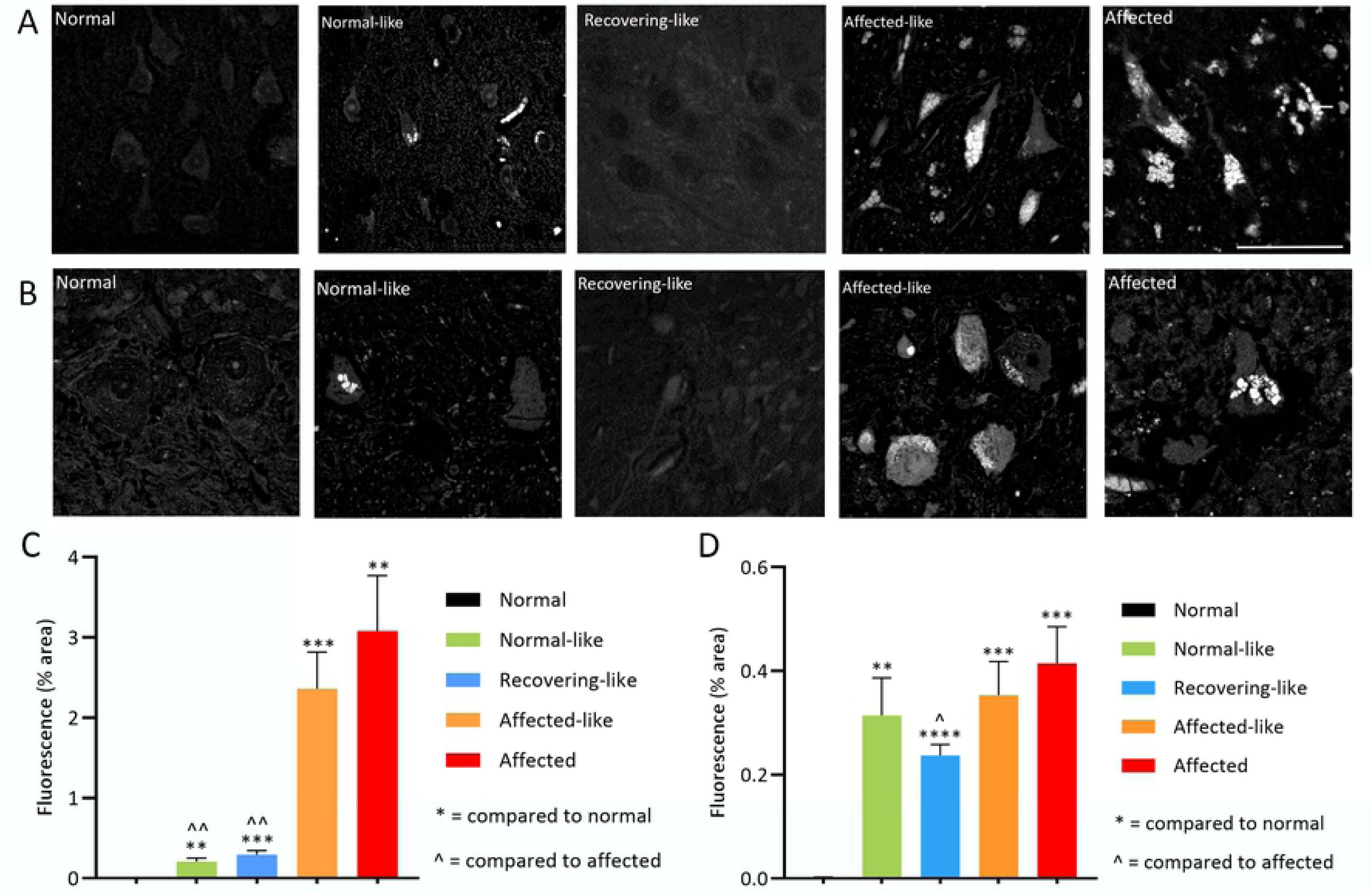
Fluorescent storage body accumulation in the cortex and cerebellum. Representative images and quantification of fluorescent storage body accumulation in the cortex (**A, C**) and cerebellum (**B, D**). Storage bodies accumulate throughout the cortex and cerebellum of affected controls and affected-like chimeras. In particular, pyramidal cells in the cortex and Purkinje cells in the cerebellum are densely packed with globular storage deposits. Conversely, storage bodies in the cells of the normal-like chimera, N1, are not tightly packed and many cells exhibit no storage at all, similar to normal controls in which storage bodies do not accumulate. No overt storage body deposits were observed in the remaining normal-like chimera, N2, and the recovering-like animals, R1, R2, R3 although quantitative analysis revealed more low-level background fluorescence in these animals than normal controls. Scale bar represents 50μm. ** and ^^ indicate P<0.01, *** indicates P<0.001, **** indicates P<0.0001.

Small storage deposits accumulated within cells of all neocortical, subcortical and cerebellar regions in one normal-like chimeric animal, N1, but at a much lower incidence than observed in affected animals, only one in 20-30 cells containing some storage bodies. Cells with globular storage deposits were present alongside cells which showed no accumulation (Fig 5). Storage in some cells resembled that in cells in affected animals, whilst others had only a few globular deposits present along the periphery of the cell perikaryon. No regional variation in accumulation was observed, all regions exhibiting the same incidence and degree of storage body accumulation. All brain regions of the normal-like chimera, N2, and the three recovering animals, R1, R2 and R3 lacked overt storage bodies although quantitative thresholding image analysis detected more nascent fluorescence in these animals than normal animals (Fig 5D).

### Astrocytic and microglial activation within the chimeric brains

Astrocytosis, revealed by GFAP immunoreactivity, was intense in the pia mater and hypertrophic astrocytes were evident across all cortical layers in the affected-like animals, A1 and A2. This immunoreactivity formed a dense meshwork, particularly prominent in upper cortical layers and was slightly less intense than that in affected controls, suggestive of a less advanced astrocytic response (Fig 6A). In contrast GFAP reactivity in the normal and recovering-like chimeras was confined to protoplasmic astrocytes consistent with that in normal control animals. Quantification of GFAP staining in the cortex revealed that affected-like animals had significantly higher levels of GFAP compared to normal control animals, but not as high as affected control animals (Fig 6B). Normal-like animals had significantly lower GFAP levels compared to affected controls (Fig 6B). No differences were observed between the GFAP staining of the subcortical or cerebellar regions of any chimeric, normal and affected control animals.

**Fig 6.**
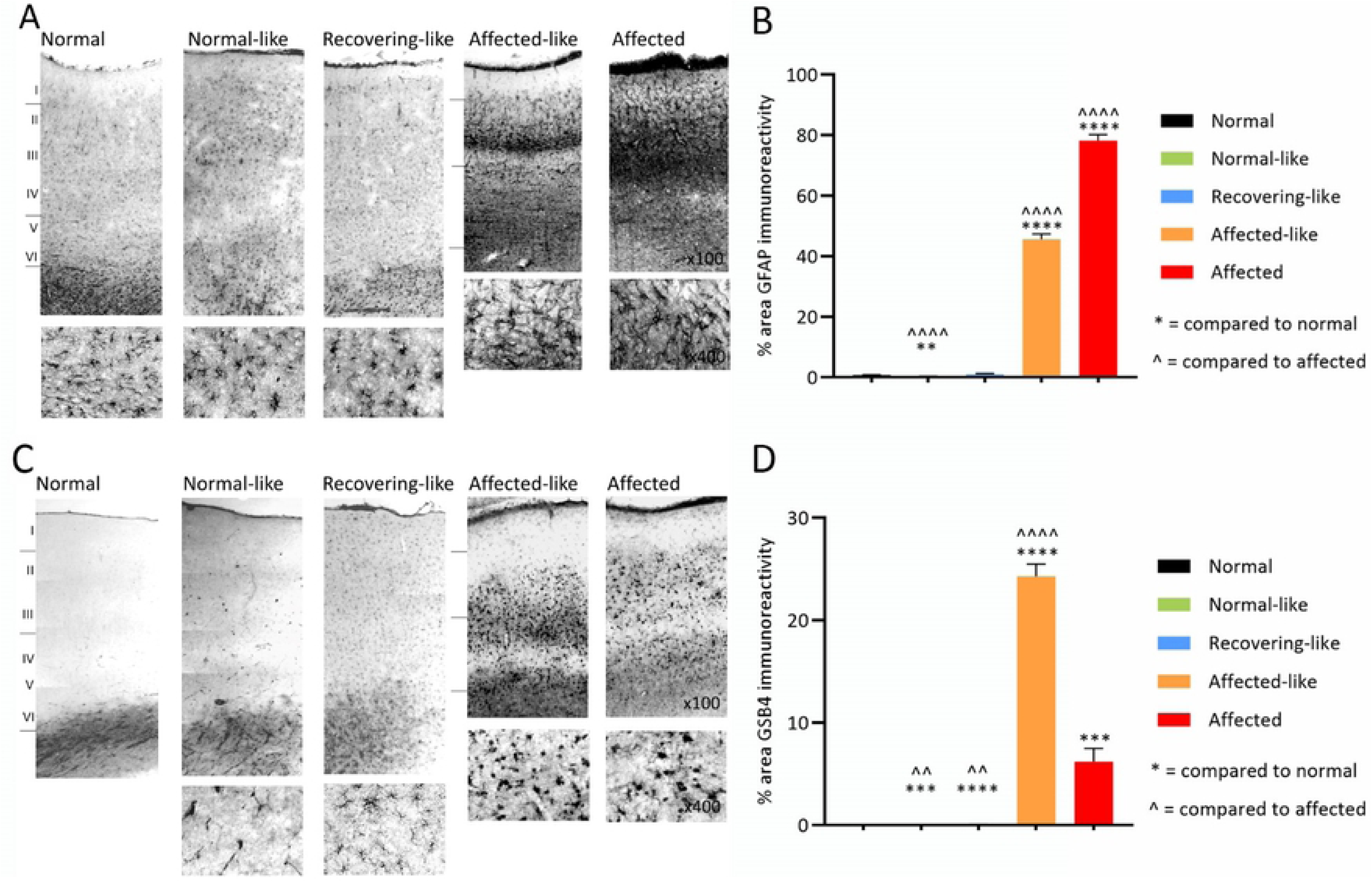
Glial activation in the occipital cortex. Representative images and quantification of astrocytic GFAP staining (**A, B**) and microglial GSB4 staining (**C, D** of the occipital cortex). Affected-like chimeras display prominent astrocytic and microglial activation consistent with age-matched affected controls. One normal-like and one recovering-like chimera had scattered activated microglia and isolated brain macrophages throughout the cortical layers, whereas the remaining normal-like and recovering-like chimeras lack glial activation. See S2 Fig for all images. ^^ indicate P<0.01, *** indicates P<0.001, **** and ^^^^ indicate P<0.0001.

Microglial activation was consistent with the GFAP findings. GSB4 staining of the two affected-like chimeras, A1 and A2, revealed an intense microglial response in cells with hypertrophied cell bodies and retracted processes typical of brain macrophages (Fig 6C, D). Staining was particularly intense in two prominent, continuous bands, upper cortical layers II-III, and lower cortical layers V-VI. The pattern and distribution of staining in these two sheep was comparable to that in affected animals and there was no regional variation in staining intensities.

GSB4 staining of the two recovering, R2 and R3, and normal-like N2 chimeras was confined to white matter capillaries, likely an artefact of prolonged immersion fixation (Fig 6C). A few flattened, elongated perivascular cells were present but no activated perivascular macrophages were detected, consistent with the lack of astrocytosis in these animals. One normal-like, N1, and a recovering chimera, R1, displayed GSB4-positive microglia scattered throughout cortical layers I-VI (Fig 6C). The majority had a ramified morphologies and small cell bodies, characteristic of resting, non-reactive microglia but occasional cells with thicker, retracted processes were scattered throughout all cortical layers, suggestive of cells transforming to activated brain macrophages. Quantification of GSB4 staining in the cortex revealed significantly higher levels of GSB4 in normal-like and recovering-like animals compared to normal controls, but these levels were still significantly lower than affected controls (Fig 6D). No regional or cortical layer differences in staining or intensity were observed.

### Extended neurogenesis in the chimeric sheep brain

Immunohistochemistry for a marker of developing and migrating neurons, polysialated neuron cell adhesion molecule (PSA-NCAM), was used to explore neurogenesis in the chimeric brains (Fig7). All seven chimeras displayed more intense PSA-NCAM immunoreactivity than normal animals, along the subventricular zone (SVZ) and within white matter tracts and cortical grey matter. There was intense staining along the SVZ in the two affected-like animals, A1 and A2, with a conspicuous band of cells and fibres even at advanced stages, a phenomenon also seen in affected controls (Fig 7A). Many individual small cell bodies were stained and larger cells with multiple processes were particularly evident in more rostral regions. Immunoreactivity in SVZs of the normal-like and recovering-like chimeric was more intense than in normal controls but less than in the SVZ of affected animals.

**Fig 7.**
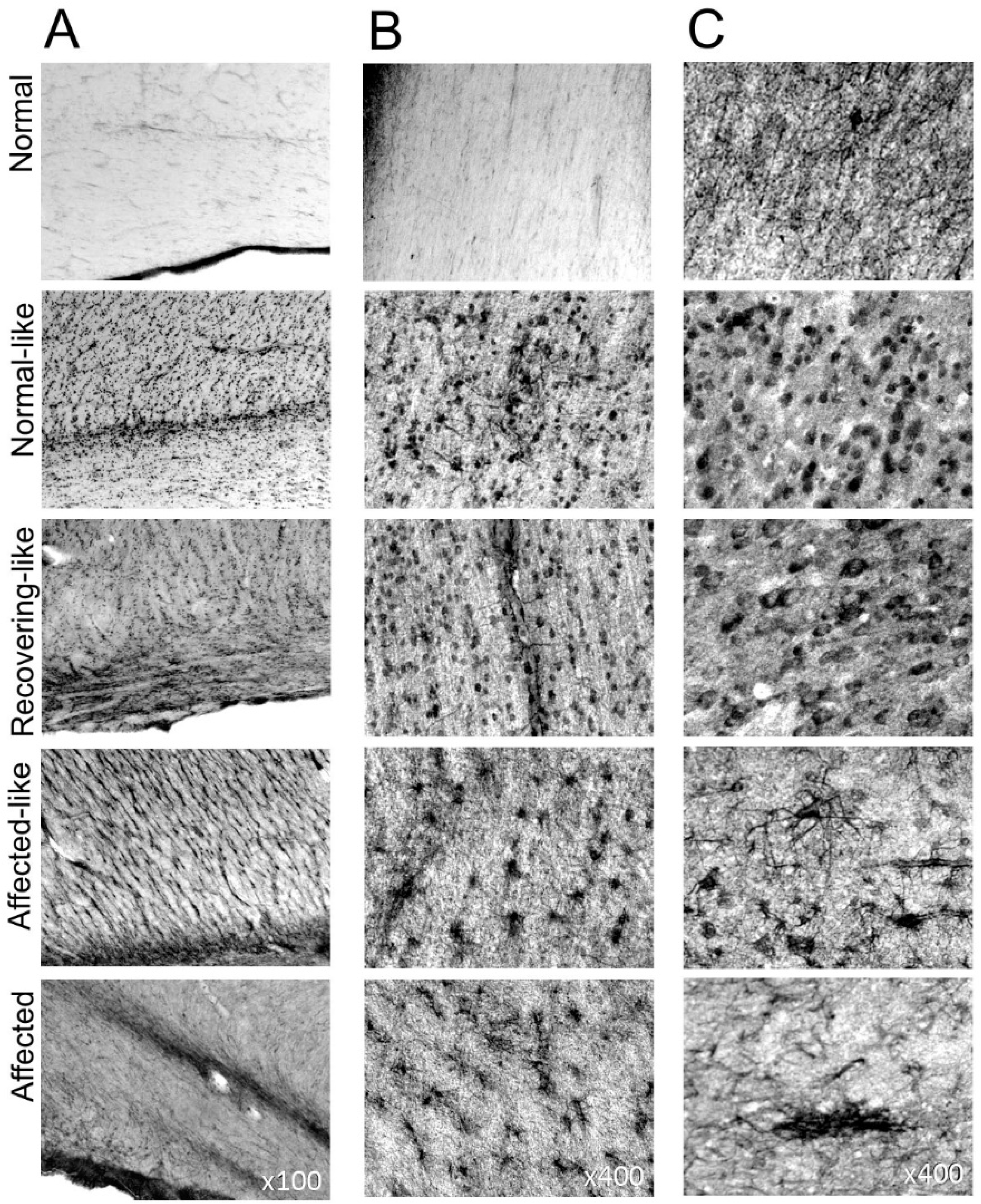
Neurogenesis in the sheep brain. Representative images of PSA-NCAM staining along the SVZ (A) and within the white matter (B) and grey matter (C) of control and chimeric animals. A band of newly generated and migratory cells are visible along the SVZ (A) in severely affected animals but not in normal controls. Staining is significantly increased along this region in all chimeric animals compared to normal controls and cells can be seen migrating along white matter tracts (B) towards the cortex (C). Large cellular aggregates are prominent within layer II of the grey matter in affected animals whereas chimeras display newly generated cells dispersed throughout all cortical layers. See S3 Fig for all images.

All seven chimeric brains contained migrating PSA-NCAM positive cells and fibres with radial orientations within white matter tracts (Fig 7B). Staining within cortical grey matter regions was intense (Fig 7C). Immunopositive cells in the cerebral cortex of all chimeras differed morphologically from those seen in the affected brains. Large cellular aggregates were seen only at the cortical layer I/II boundary in affected animals. In contrast, cells in the affected-like animals, A1 and A2, had intensely stained perikaryon and multiple dendritic processes, present at a high incidence throughout all cortical layers, particularly the upper layers, but they did not cluster. The morphology of the PSA-NCAM stained cells in the chimeras N1, N2, R1, R2 and R3 was different again. Numerous cell bodies, with occasional apical dendrites, were immunostained uniformly across the cortical layers in all these recovering and normal-like animals. These were absent or had a very different morphology in the normal controls. PSA-NCAM positive cells were also detected within the dentate gyrus of the hippocampus but no qualitative differences in the staining intensity was noted between affected and normal controls or chimeric animals in this region.

### Retinal pathology in chimeric eyes

Retinal thickness and retinal fluorescent storage body accumulation was assessed. Retinas of normal-like and recovering-like chimeras had healthy looking layer morphology and were significantly thicker than those of affected controls (Fig 8A, B). The thickness of the normal-like retinas was not significantly different from normal controls, the recovering-like retinas were slightly thinner and affected-like retinas were much thinner and had disrupted layer morphology. Cell loss was evident from both affected-like and affected control retinas, primarily from the outer nuclear and photoreceptor layers (Fig 8A).

**Fig 8.**
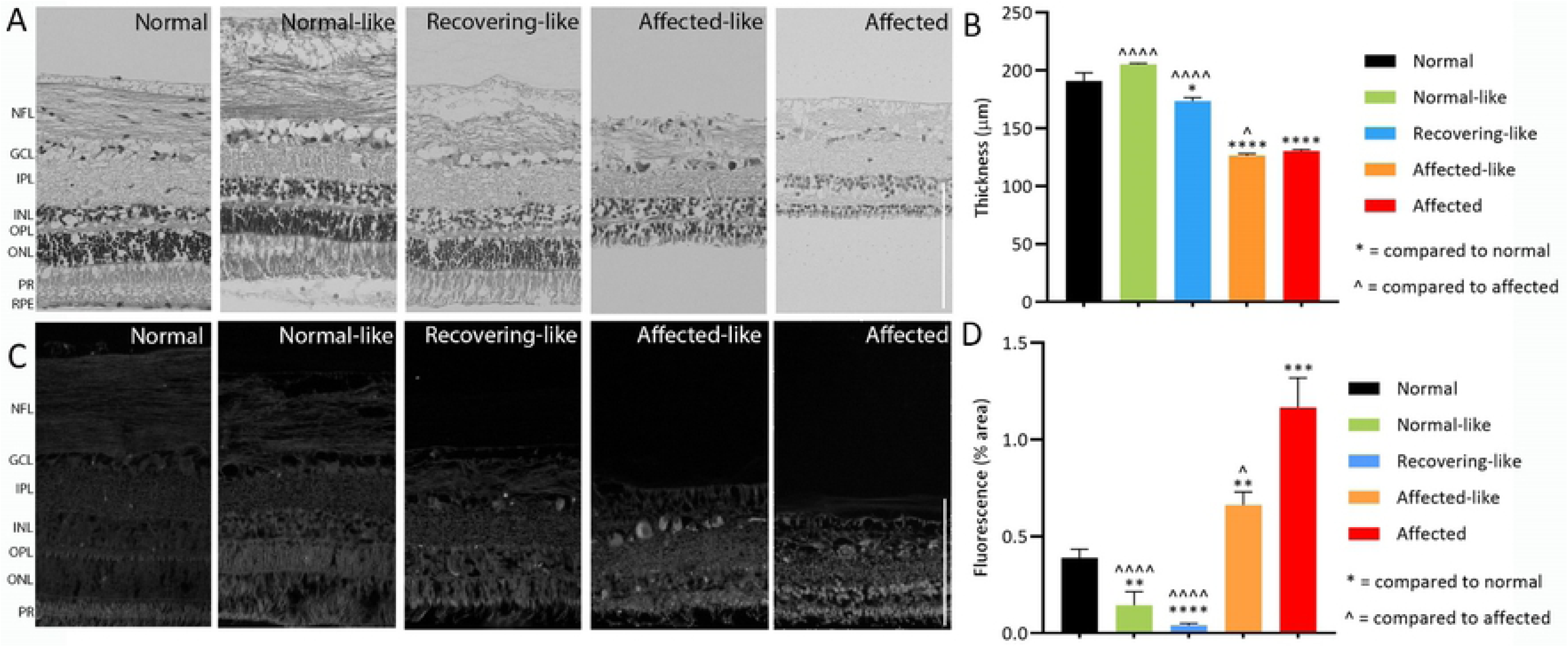
Retinal thickness and fluorescent storage body accumulation. Retinal thicknesses (**A, B**) in normal-like and recovering-like chimeric sheep retina were similar to those of normal controls, while affected and affected-like chimeras had significantly thinner retina. Fluorescent storage body accumulation (**C, D**) was highest in affected controls, although affected-like chimeras still had significantly more than normal controls. NFL: nerve fibre layer; GCL: ganglion cell layer; IPL: inner plexiform layer; INL: inner nuclear layer; OPL: outer plexiform layer; ONL: outer nuclear layer; PR: photoreceptor layer; RPE: retinal pigment epithelium. * and ^ indicate P<0.05, ** indicates P<0.01, *** indicates P<0.001, **** and ^^^^^ indicate P<0.0001. Scale bars represent 100µm.

Fluorescent storage bodies were most prominent in the retinas of affected-like chimeras, particularly in the ganglion cells and inner and outer nuclear layers (Fig 8C). Quantification of storage material revealed that although levels of storage were significantly higher than normal controls, they were also slightly lower compared to affected controls. Normal-like and recovering-like chimeras had significantly lower retinal fluorescence than affected controls and even less than the endogenous level seen in normal retinas (Fig 8D).

## Discussion

As with other lysosomal storage diseases, the NCLs caused by soluble enzyme deficiencies are more likely to benefit from cross-correction than are diseases caused by intracellularly contained membrane bound proteins. Exogenous soluble enzymes can be provided by enzyme replacement, stem cell or gene therapies, as has been achieved in animal models and human patients [33–37].

Defects in membrane-bound proteins, including CLN6, an endoplasmic reticulum resident protein of unknown function [14–16], are considered harder therapeutic targets as the disease mechanism is anticipated to be intracellular. However, intracerebroventricular delivery of CLN6 gene therapy to the neonatal *Cln6* mutant mouse has been shown to prevent or drastically reduce all the pathological hallmarks of NCL, as well as improving behaviour and extending survival [38]. Safety and CNS targeting was confirmed in non-human primates [38] and a phase I/II CLN6 gene therapy trial is underway (Clinical trial.gov identifier: NCT02725580). However, brain size and complexity are important and it is not at all clear if the rescue of some cells can rescue the phenotype of a larger complex brain where cells are less accessible to transfection.

In ovine CLN6, there is widespread storage body accumulation throughout the cells of the body [39] implying that the underlying pathological insult must be similar in all cells, yet severe degeneration is confined to the CNS and is regionally defined [13]. Cellular location and interconnectivity, rather than phenotype, are considered major determinants of neuron survival [12], indicating that intercellular interactions may be possible. Having normal and CLN6 affected cells intermixed within tissues, as in the chimeric sheep here, is a direct way to establish whether neuronal cells expressing the CLN6 protein influence other non-expressing neuronal cells and test the argument that therapies based on cellular cross-correction are realistic for CLN6 NCL in a complex large animal brain.

### Development and heterogeneity of chimeric animals

The considerable variation in the proportions of normal to affected cells in tissues in the seven chimeric animals highlighted the extent of heterogeneity of animals constructed by embryo aggregation, consistent with previous findings that the relative colonization by genotypically different cell lineages in chimeric animals can differ between tissues and animals [40, 41]. If normal and affected cell mixing occurs prior to formation of the inner cell mass (ICM) both cell types contribute to the ICM and its subsequent differentiation into the ectodermal, mesodermal and endodermal layers, and thus to all the tissues of the body. Furthermore, the variation of cell proportions within the body is dependent on the extent of mixing and proportion of each cell genotype in the ICM. For the affected-like chimeras, cells with an affected genotype contributed most to formation of the ICM, whereas the normal and recovering-like chimeras had varying proportions of both normal and affected cells in most tissues, as has been observed previously [42–44]. Regional variations were also observed within brain tissues in most of the chimeras, as noted in previous studies [24, 25].

A summary of results from this study is presented in Table 2. The affected-like animals, A1 and A2, displayed increased neurogenesis as indicated by PSA-NCAM staining, intense glial activation, prominent storage body accumulation and severe neurodegeneration within all cortical brain regions, similar to that in affected controls and in line with previous results [6,11–13,45–47]. Consistent with these findings was a progressive loss of vision, intracranial volume and cortical thickness. Nevertheless PSA-NCAM staining showed that the presence of normal cells had a profound effect in these animals. Immunoreactive cells were not confined to cellular aggregates as seen in affected controls but were found throughout all cortical layers. GFAP staining also revealed a less advanced astrocytic response than that in affected controls. However, GSB4 histochemistry displayed a comparable pattern and distribution of staining to that in affected animals.

**Table 2.** Phenotypic, genotypic and histological status of chimeric animals.

| Classification | Animal | Endpoint (months) | Terminal weight (kg) | Phenotype |  | Genotype (% affected)<sup>b</sup> |  |  | Storage | Gliosis | Neurogenesis | ICV (% normal)<sup>c</sup> |
| --- | --- | --- | --- | --- | --- | --- | --- | --- | --- | --- | --- | --- |
|  |  |  |  | Coat pattern<sup>a</sup> | Vision | Blood | CNS | Non-CNS |  |  |  |  |
| Affected-like | A1 | 25.5 | 43.4 | Affected | Blind | 100 | 100 | 12.5-100 | Typical | Yes | Yes | 73 |
|  | A2 | 19.5 | 37.0 | Affected | Blind | 100 | 100 | 100 | Typical | Yes | Yes | 80 |
| Recovering-like | R1 | 41 | 94.5 | Normal | Normal | 3 | 3-25 | 0-50 | None | Yes | Yes | 78 |
|  | R2 | 26 | 74.0 | Normal | Normal | 0 | 3-6 | 0-3 | None | No | Yes | 87 |
|  | R3 | 26 | 70.2 | Normal | Normal | 12.5 | 6-75 | 6-50 | None | No | Yes | 100 |
| Normal-like | N1 | 26 | 86.8 | Normal | Normal | 0 | 12.5-50 | 12.5-87.5 | Some atypical | No | Yes | 103 |
|  | N2 | 26 | 71.0 | Normal | Normal | 50 | 3-25 | 6-75 | None | No | Yes | 107 |

<sup>a</sup>Predominant coat pattern was either affected (South Hampshire) or normal (Coopworth)
<sup>b</sup>Percentage of CLN6 affected DNA in blood, central nervous system (CNS) and non-CNS tissues estimated
<sup>c</sup>ICV (intracranial volume) was determined as a percentage of normal

Histological analysis of the retinas of chimeric animals revealed pathology in keeping with their normal-, recovering-, or affected-like classifications. The retina of CLN6 affected sheep exhibits severe atrophy of the photoreceptor layer, and the outer nuclear and plexiform layers [46,48,49]. Accumulation of lysosomal storage, particularly in the ganglion cell layer is another common feature of the retina in ovine NCL [48,50–52]. Affected-like chimeric animals showed levels of atrophy and lysosomal storage similar to that in affected controls, normal-like chimeras showed no signs of retinal atrophy and very little storage, whilst recovering-like animals sat somewhere in between. These differences in retinal pathology indicate that CLN6-defcient cells have an influence over normal cells, and vice versa.

### Normal- and recovering-like chimeric animals

Intercellular communication affecting pathology was evident at both the gross and histological level in the normal-like and recovering-like chimeras. The normal-like chimeras, N1 and N2, with genotypes indicative of a more balanced presence of normal and affected cells within tissues, displayed a lack of glial activation even at advanced ages. Similarly, storage body accumulation was only evident in some cells in one animal, N1, and the extent of storage in these cells was minor compared to storage in cells of affected animals at younger ages. These findings indicate cross-cell correction of affected cells by normal cells resulting in a reduction or absence of storage bodies, removal of them, or a halt in the process of accumulation. This phenomenon is also shared in the other normal-like and recovering-like chimeras, there being no storage accumulation observable at the time of sacrifice. Intracranial volume and cortical thickness findings were all consistent with normal controls and there was no loss of vision.

Analysis of the recovering chimeras, R1, R2 and R3, indicated a larger proportion of normal cells than affected cells in most brain regions but some dramatic variations were evident in animal R3 (Table 2). Although intracranial volumes of all three of these animals were below normal, those of animals R2 and R3 had recovered to about normal volume by two years of age and R1 appeared to have a progressively recovering brain volume. All three animals had normal cortical thickness measurements, none went blind and there was no evidence of storage body accumulation (Table 2). Glial activation, observed only in animal R1, was less advanced than in affected controls, there being fewer activated cells stained and it should be noted that this animal was euthanised at 41 months of age, so may have experienced some typical age-related gliosis.

In stark contrast to the lack of neuroblastic activity in the normal brains, extended neurogenesis was evident in the brains of all the chimeric sheep and was particularly robust in the normal and affected-like chimeras. PSA-NCAM positive cells were confined to large cellular aggregates in upper cortical layers of the affected-like chimeric brains and there were immunopositive neurons throughout all cortical layers of the normal-like and recovering-like brains. As revealed by Nissl staining, these normal and recovering chimeras had an intact laminar distribution of cells and normal control cortical thickness measurements indicating a lack of neurodegeneration, whereas affected sheep displayed a laminar reorganisation corresponding to the occurrence of disease symptoms [6,9,13]. It is possible that the newly generated cells originating from normal neural progenitor cells (NPCs) undergo successful migration and distribute correctly throughout all cortical layers in these animals. The lack of glial activation and inflammatory response in the normal-like and all but one (R1) recovering-like chimeras indicate that the newly generated cells are being borne into a microenvironment conducive to cell maturation and survival. These findings are reflected in the intracranial volume data and suggest that migration of corrected cells, in combination with a neurotrophic environment, result in newly generated cell survival leading to recovering intracranial volumes and disease amelioration.

As all these animals are chimeras, it is probable that some NPCs will be of an affected CLN6 genotype and some of a normal genotype. Affected degenerating cells could theoretically be replaced with functional, unaffected cells or with cells carrying the *CLN6* mutation. Therefore, replacement of affected cells by normal, unmutated cells may occur at a slower rate in some animals and in some brain regions depending on the population of progenitor cells from which the new cells arise or alternatively, if glial activation is ablated, mutated cells may survive for an extended period in the absence of a detrimental inflammatory environment. The three animals that initially had lower intracranial volumes, R1, R2 and R3, had an apparent high colonisation of normal cells in the brain, as inferred by histological and genotypic analysis. Their reduced intracranial volume may have been a consequence of early loss of affected neurons and subsequent progressive replacement by neuroblasts generated from normal NPCs. Animal R1 also displayed microglial activation, albeit at a much lower intensity than affected animals. This less intense glial activation may have caused neuronal death but at a slower rate than in affected animals, hence enabling a progressive rate of cell replacement.

Aggregation chimera production provides both normal and affected cells to the prenatal brain and normal cells may have inhibited the early glial response proposed to be a causative factor in neurodegeneration and pathology in the ovine CLN6 model [11, 13]. The initial affected:normal cell proportions, and changes in these proportions over time, are not known for all these animals. Location and connectivity, not phenotype, determine neuronal survival in ovine CLN6 [12] and it could be that there are critical cell types and brain regions required for normal development, accounting for the differential developmental pathways in these chimeras.

### Cross-cell communication and neurotrophic factors

This chimera study strongly indicates that although CLN6 is a membrane bound protein the consequent defect is not cell intrinsic. The fact that normal cells appeared to alter the fate of affected cells in the normal-like and recovering-like chimeras suggests that it may be involved in the processing of secreted factors, which when released provide a specific survival or anti-apoptotic signal to affected cells or create a better growth environment able to support CLN6-deficient cells. Although the critical threshold of normal cells required to bring about therapeutic benefit is unknown, it is clear that not all cells need to be corrected.

The proposed role of CLN6 in pre-lysosomal vesicular transport [15] suggests that the sorting and processing of factors like neurotrophins and their receptors could be affected in ovine CLN6, resulting in their reduced expression. Neurotrophic factors promote neuronal survival, stimulate axonal growth and play a key role in construction of the normal synaptic network during development [53]. In adulthood, they help to maintain neural functions, therefore any alterations in their local synthesis, transport or signalling could adversely affect neuronal survival and lead to neuronal death [54, 55]. A number of studies have shown that a loss of neurotrophic support for selective neuronal populations may contribute to the pathology of other neurodegenerative diseases including Parkinson’s, Alzheimer’s and Huntington diseases [55, 56]. In some circumstances, treatment with neurotrophic factors including nerve growth factor (NGF), brain-derived neurotrophic factor (BDNF), glial cell-derived neurotrophic factor (GDNF), insulin-like growth factor (IGF) and neurotrophins 3 and 4/5 can prevent cell loss.

Another possibility is that the CLN6 protein influences the lysosomal targeting, sorting and secretion of one or more soluble lysosomal proteins as does CLN8, another membrane bound NCL protein [57]. In this case the return of function to the affected cells in the chimeras would arise from cross-correction with a soluble factor secreted from normal cells.

Targeting of progenitor cells in the SVZ which give rise to neuroblasts that migrate to regions of neurodegeneration, working in concert with cross-correction, could extend the zone of therapeutic benefit and gene therapy studies using these techniques in the ovine CLN6 model has been reported [58] and are ongoing. Such results should prompt similar investigations in other NCLs resulting from presumed membrane bound protein defects. They also indicate a good prognosis for intracerebroventricular delivery of CLN6 gene therapy to large complex brains as it is not necessary to transduce all the relevant cells.

## Supporting information

**S1 Fig. Nissl staining of the occipital cortex in each individual chimeric sheep, compared to normal and affected controls.**

**S2 Fig. Neuroinflammation in each individual chimeric sheep. A)** GFAP astrocytic staining and **B)** GSB4 microglial staining animals of normal, affected and chimeric occipital cortex.

**S3 Fig. Neurogenesis in each individual chimeric sheep.** PSA-NCAM staining along the SVZ (A) and within the white (B) and grey matter (C) of control and chimeric animals.

